# An extended Stokes’ theorem for spiral paths: applications to rotational flows in *Trachelospermum jasminoides* stems and flowers

**DOI:** 10.1101/2025.01.15.633128

**Authors:** Arturo Tozzi

**Author notes:** (corresponding author) 1155 Union Circle, #311427 Denton, TX 76203-5017 USA.

## Abstract

The traditional Stokes’ theorem connects the macroscopic circulation along a closed boundary to the microscopic circulation across the surface it encloses. However, it proves inadequate for addressing complex geometries such as helicoidal paths, non-planar flow patterns and dynamic systems with open boundaries. We introduce an extension of Stokes’ theorem (EST) that provides a robust tool for interdisciplinary research in spiral/helicoidal dynamics, facilitating the evaluation of rotational forces and circulation in both natural and engineered systems with open boundaries. We apply EST to model the rotational dynamics of flower petals and the helical forces within the stems of *Trachelospermum jasminoides*, known as star jasmine. For the flower, we demonstrate the equivalence between the line integral along the petal boundary and the surface integral over the enclosed disk, effectively capturing the uniform rotational stress generated by tangential forces. EST enables the analysis of external factors such as wind or pollinator interactions, while providing valuable insights to deepen our understanding of floral mechanics and petal growth patterns. For the stem, linking microscopic circulatory forces to macroscopic flow patterns, we demonstrate the interaction of torsional and bending stresses caused by the helical geometry. This finding has significant implications for understanding plant growth biomechanics and structural stability as well as for quantifying nutrient and water transport within stems, where spiral dynamics play a pivotal role. In summary, EST streamlines the analysis of rotational and translational forces in systems governed by spiral and helicoidal dynamics, including physical and biological phenomena such as phyllotaxis and plant growth.

## INTRODUCTION

Stokes’ theorem (henceforward ST) is a fundamental principle of vector calculus that bridges the macroscopic circulation along a closed boundary with the microscopic circulation over the enclosed surface (Green, 1828; Schey, 1997). Extending the principles of Green’s theorem (GT) which applies to two-dimensional regions, ST provides a powerful framework for analyzing flows and circulations in three-dimensional spaces, uncovering profound connections between the local properties of vector fields and their global behaviour. GT and ST are effective tools for solving problems related to physical closed systems with clearly defined boundaries such as airflow circulation around wings, electromagnetic fields in circuits, surface heat flux, Coriolis-driven hemispherical flows, Earth’s deep interior dynamics (Craven, 1964; Arfken, 1985; De Villiers, 2006; Livermore et al., 2013; Snieder, 2015; Aubert and Finlay, 2019; Vines et al., 2021; Yang et al., 2023). In biology, the two theorems contribute to understanding blood flow, electrical activity of the brain, growth patterns in ecosystems (Tozzi and Peters, 2023; Bressan et al., 2022). Yet, the classical ST is inherently limited to surfaces and boundaries that are closed, leaving a significant gap in its applicability to open, non-planar geometries. Indeed, many natural and engineered systems exhibit spiral or helical dynamics where forces and flows do not conform to closed loops or planar surfaces but rather are characterized by open, three-dimensional trajectories. Examples include the helical paths of tornadoes, magnetic vortices and spiral galaxies as well as bacterial motility and phyllotaxis of plants (Blaser et al., 2024; Sachkou et al., 2019; Reinhardt and Gola, 2022).

The novelty of this work lies in extending ST to accommodate spiral flows and helicoidal paths. By linking macroscopic and microscopic circulation properties, the extended theorem simplifies the evaluation of forces in systems with open, three-dimensional geometries. We utilize EST to analyse two biological scenarios: 1) the rotational forces in spiral flower petals and 2) the torsional stresses in helical plant stems, both exemplified by *Trachelospermum jasminoides*, commonly known as star jasmine. For the flower, EST captures the uniform rotational stresses induced by tangential forces acting along a circular boundary. This is achieved by demonstrating the equivalence of the line integral along the petal boundary and the surface integral over the enclosed disk. For the stem, EST quantifies the interaction between bending and torsional stresses caused by the helical geometry.

This paper is structured as follows. First, we present the mathematical treatment of EST, including its derivation and parameterization for helicoidal paths. Next, we validate the theorem using the specific example of *Trachelospermum jasminoides’* flowers and stems. Finally, we discuss the broader implications of EST, highlighting its potential to unify the study of dynamical systems with open boundaries.

## MATERIALS AND METHODS

This study is grounded in a generalized form of Stokes’ theorem, adapted for spiral flows, which facilitates the analysis of forces and circulation in systems with helicoidal or spiral geometries. We aim to prove that, given a continuously differentiable, orientable helicoidal spiral vector field, the macroscopic circulation represented by the integral of a differential form over its surface equals the microscopic circulation represented by the volume integral of the curl perpendicular to the surface. The main challenge here is in defining the notion of a boundary in case of an open helicoidal spiral path, moving beyond the classical case of paths evaluable through ST.

Stokes’ Theorem (ST) from vector calculus relates the surface integral of the curl of a vector field over a surface **S** to the line integral of the vector field along the boundary curve ∂S of the surface (**Figure 1**). In its general form, ST asserts that

**Figure 1.**
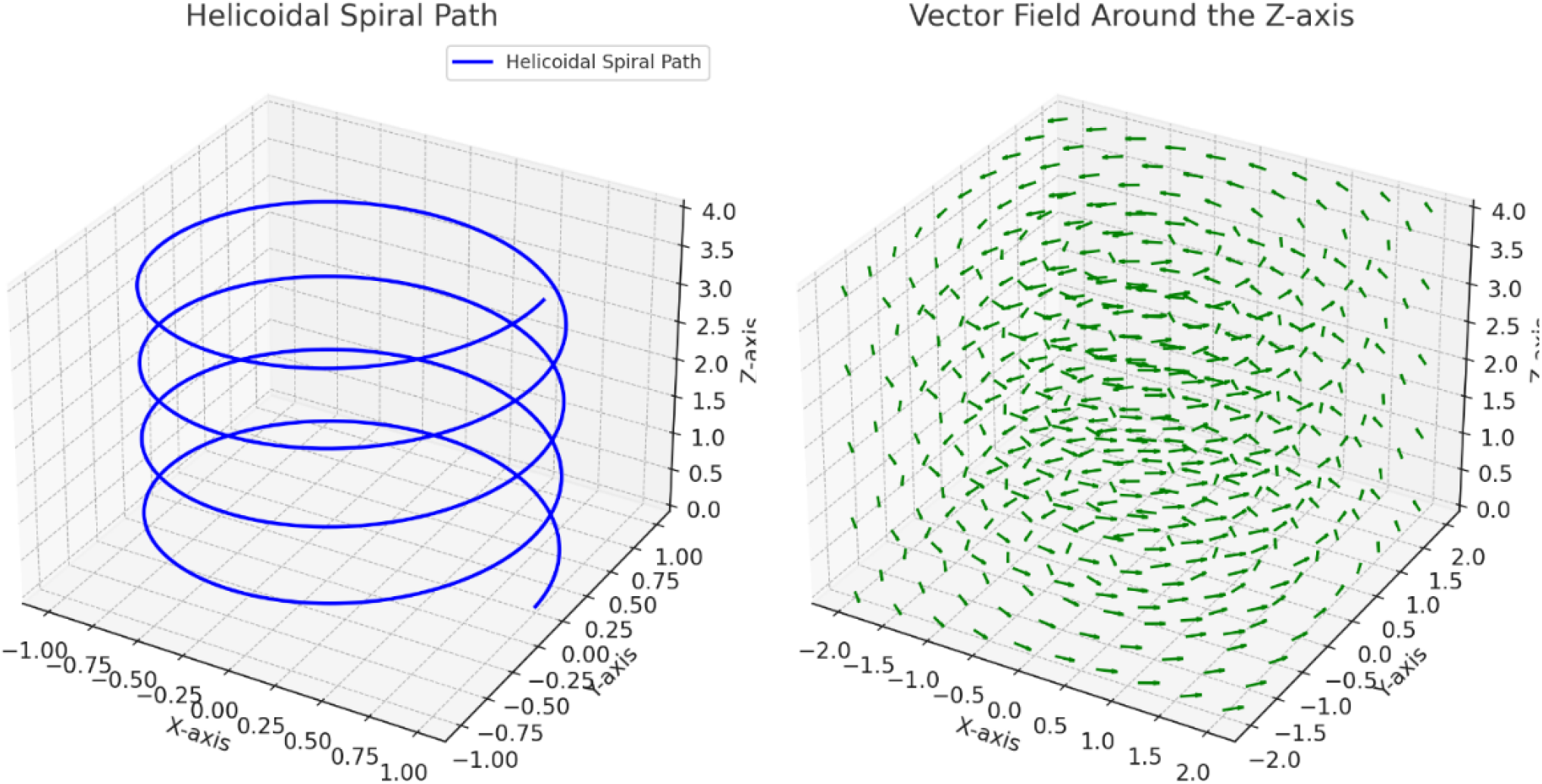
Diagrams depicting a helicoidal spiral (**left**) and the behavior of a vector field (**right**) around the z-axis. The left diagram illustrates a helicoidal spiral path, showcasing the interplay of rotational and translational motion along the z-axis. The right diagram represents a vector field with circular flow centered around the z-axis.

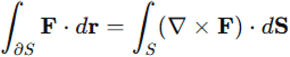

where **F** is a a continuously differentiable two-dimensional vector field, ∂S is the closed boundary curve of the surface **S** that can be bended and stretched, d**r** is a differential element of the curve, d**S** is the differential element of the surface area, and ∇ ⨯ **F** is the curl of the vector field, i.e., a vector operator characterizing the infinitesimal circulation of vector fields in three-dimensional spaces.

ST turns line integrals of a form over a boundary into more straight-forward double integrals over the bounded region, regardless of the position of vector singularities (Zenisek 1999). For ST to apply, the normal vector representing the surface must be positively oriented (i.e., counterclockwise) with respect to the tangent vector representing the orientation of the boundary.

### Extended Stokes’ theorem (EST)

Consider a vector field **F** defined over a region in three-dimensional space. Let the surface *S* be a portion of a plane or a more general surface that is bounded by a spiral curve *γ*(*t*) The goal is to use EST to evaluate the line integral over the spiral path in terms of the surface integral of the curl of **F** (**Figure 1**).

Let the spiral curve *γ*(*t*), with *t* ∈ [*a, b*], be parameterized as

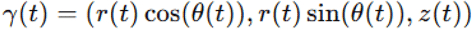

where *r*(*t*), *θ* (*t*) and *z* (*t*) describe respectively the radial, angular and vertical components of the spiral. Let’s assume that *γ*(*t*) lies on a flat plane, say the *xy*-plane, so the spiral path can be simplified to

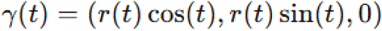

where *r*(*t*) increases as the <u>angle</u>*<u>t</u>*<u>increases.</u>

When the surface **S** is a surface spanned by the curve *γ*(*t*), **S** stands for a portion of the plane or surface generated by the spiral curve (**Figure 1, left**).

We are interested in computing the line integral of a vector field **F** along the spiral path (**Figure 1, right**). By ST, this line integral can be transformed into a surface integral involving the curl of **F**

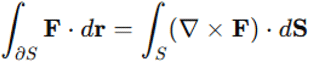

The line integral over the spiral path is:

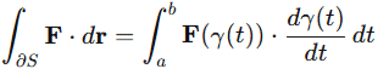

where 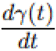 is the tangent vector to the spiral path at each point *t*.

The surface integral involves the curl of **F**, given by ∇ ⨯ **F**, and the surface normal vector associated 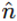 with **S**

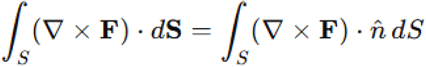

The normal vector 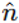 depends on the orientation of the surface, while *d*S is the differential area element of the surface. Upon achieving the extended formulation of ST, we will proceed in the next paragraphs with a detailed case study.

### Simulated case study: analyzing rotational flows in stems and flowers

EST enables the analysis of rotational and translational forces in complex systems, providing a powerful framework for exploring biological and physical dynamics. To illustrate the new theorem, we will now explore a concrete example. We will consider *Trachelospermum jasminoides*, commonly known as star jasmine, belonging to the family *Apocynaceae*. Like many climbing plants, *Trachelospermum jasminoides* displays a counterclockwise helical movement of its stems as it climbs and twines around supports, also referred to as circumnutation (Darwin, 1875; Pansanit and Pripdeevech, 2014; Canher et al., 2022) (**Figure 2**). The flowers also exhibit subtle rotational dynamics, although these movements are not as pronounced as the helical twisting of the stems (Stefanatou et al., 2025). The petals of the flowers are arranged in a spiral configuration and unfurl in a counterclockwise direction during blooming.

**Figure 2.**
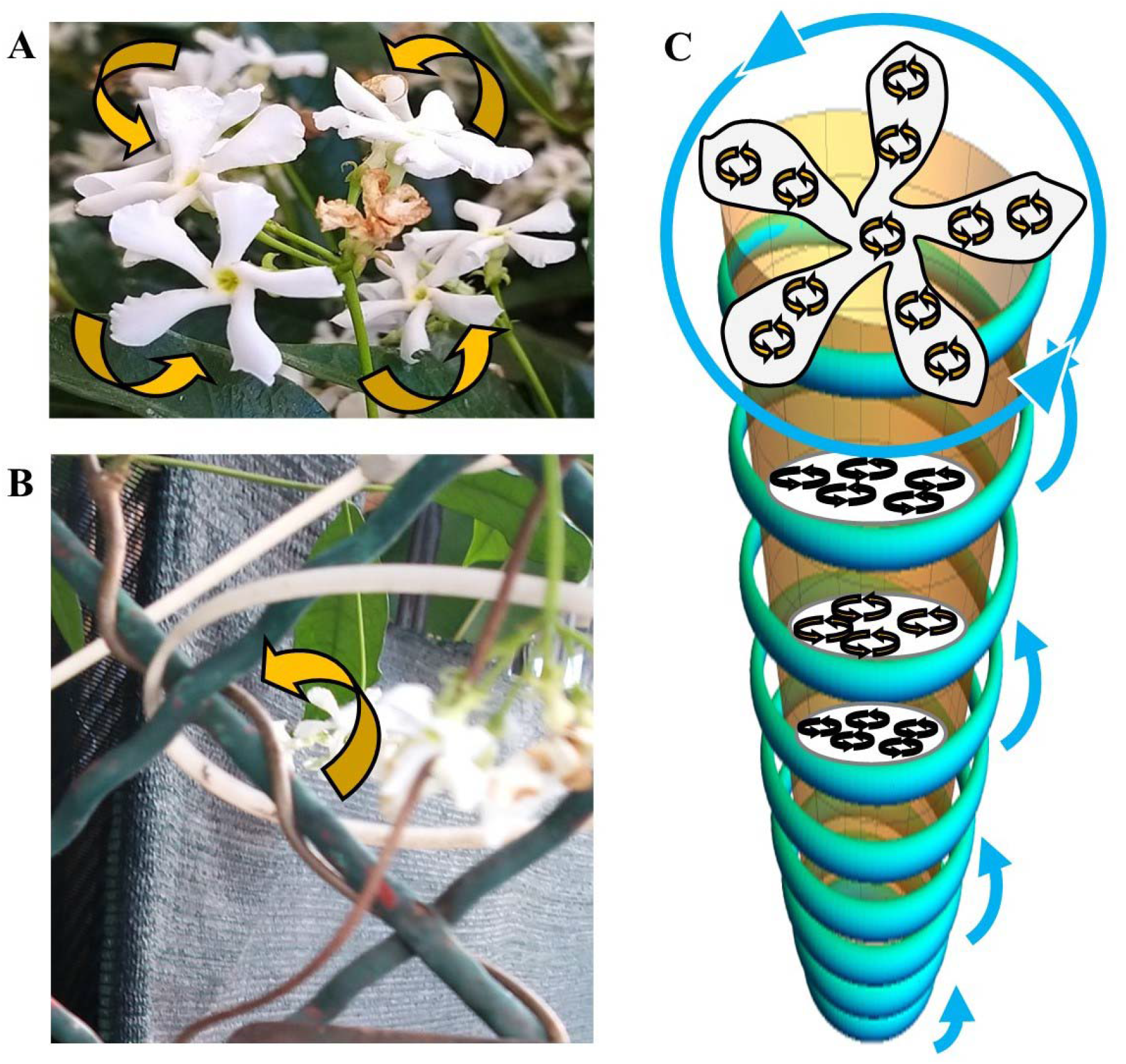
*Trachelospermum jasminoides*. The flower petals (**Figure 2A**) and the stem (**Figure 2B**) display a counterclockwise path. **Figure 2C** illustrates the geometry of the boundaries, the associated vector fields and the internal flows within the flower petals and the stem.

**Figure 3.**
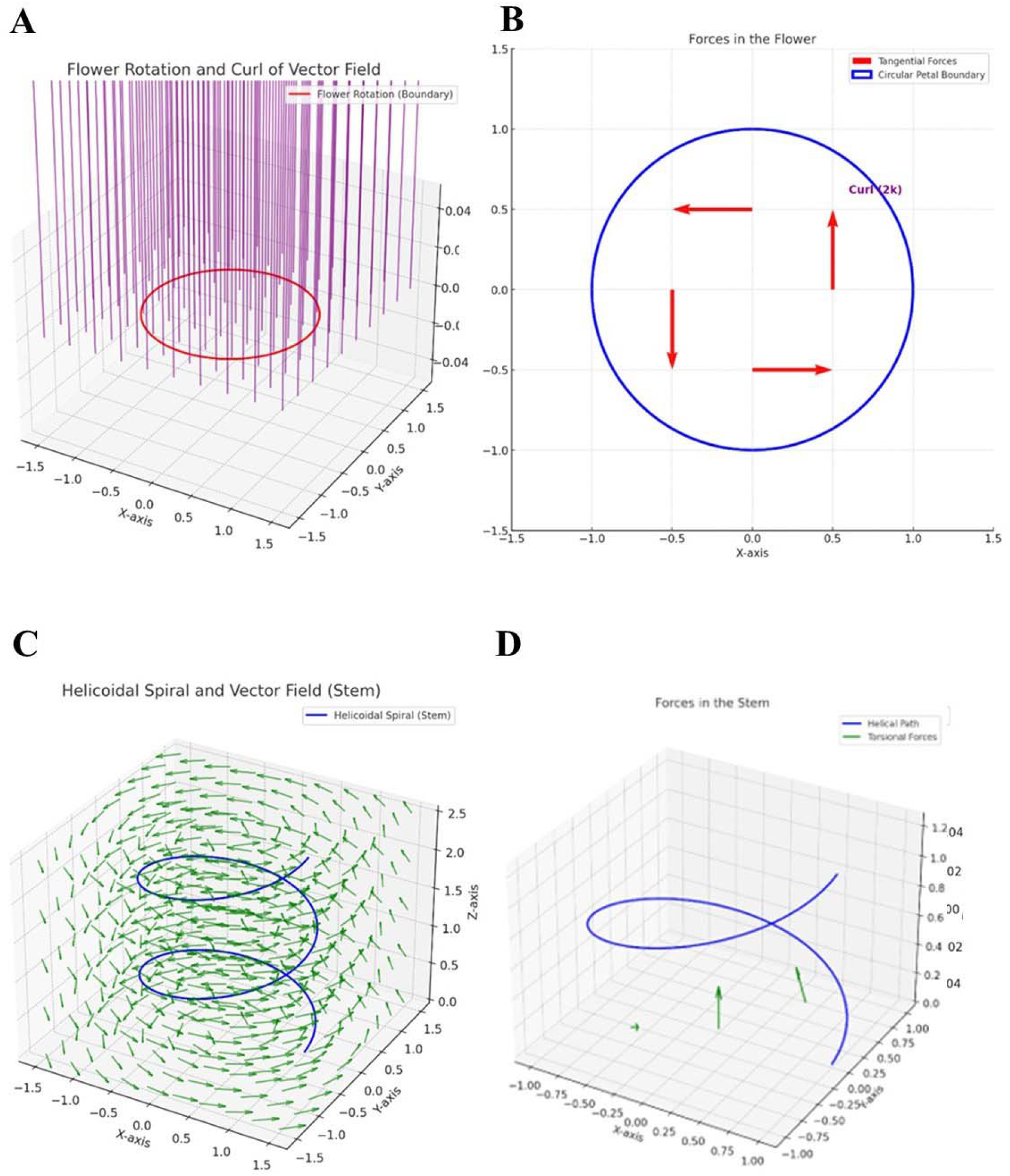
Application of the extended Stoke’s theorem to flowers (**Figures 3A-B**) and stems (**Figures 3C-D**) of *Trachelospermum jasminoides*. **Figure 3A**. Diagram illustrating the flower rotation and the curl of vector field. The red circle represents the boundary of the flower petals modeled as a planar region in the z=0 plane. The purple arrows visualize the curl of the vector field, representing the microscopic circulation that contributes to the macroscopic rotation of the flower petals. **Figure 3B**. Diagram illustrating the forces acting on the flower. The circular boundary (blue) represents the edge of the flower petals. Tangential forces (red arrows) act along the edges of the petals, showcasing the influence of external or internal factors. The calculated curl of the vector field is constant at ∇ ×F=2k (annotated in purple) in the z-direction, indicating uniform rotational stresses throughout the petal boundary. **Figure 3C**. Diagram illustrating the helicoidal spiral of the stem and the vector field. The blue curve depicts the helicoidal path of the stem, while the green arrows represent a circular vector field around the z-axis, illustrating the rotational and translational flow and its interaction with the spiral geometry. **Figure 3D**. Diagram illustrating the forces acting on the stem. The helical path (blue curve) represents the stem’s geometry. The torsional forces (green arrows), resulting from a combination of bending and twisting actions, act along the helical structure contributing to internal stress distribution. The curl vector field displays non-uniform rotational stresses with components ∇ ×F=(k,k,2k).

In our simulation, the dynamics of flower petals are modeled using a circular boundary with a radius of 0.05 m, representing the petals of a flower. Tangential forces along this boundary are applied and the resulting rotational stresses are analyzed through EST. The stem is modeled as a helicoidal path with a radius R=0.05 m and a vertical rise per turn of c=0.2 m. To evaluate the counterclockwise rotation of the flower and stem using EST, the rotation of the petals can be represented by a circular vector field, whereas the helical motion of the stem can be modelled using a helical vector field. The next step is to parameterize the flower and the stem (**Figure 2C**).

1. For the flower, a circular boundary in the plane of the petals is defined, representing the region of interest for macroscopic rotation.
2. For the stem, the helical path is parameterized using equations for a helicoidal spiral, where *x*(*t*) = *r* cos(*t*), *y*(*t*) = *r* sin(*t*) and *z* (*t*) = *ct*. Here r represents the radius, c the rise per turn and t the parameter along the path.

Subsequently, the surface S is defined for each component.

1. For the flower, the surface is a disk enclosed by the petals’ rotational motion within their plane,
2. whereas for the stem, the surface corresponds to the area traced by the helical path (**Figure 2C**).

Calculating the forces acting on the flower and stem requires applying mechanical principles that account for both internal and external forces influencing their dynamics (Smyth 2016; Loshchilov et al., 2021). For the flower’s petals, the primary force is torque or rotational force, while the stem experiences a combination of bending forces and axial torsion due to its helical structure. We will calculate these forces systematically, step by step, starting from the external forces.

### External forces acting of the flower petals and the stem

1. The rotation of the flower petals can be modeled as a torque induced by external forces such as wind, gravitational pull, biological growth forces (Tipler, 2004). Torque (*τ*) on the petals is given by:

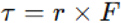

Where r is the radial vector from the center of the flower to the tip of a petal and F is the tangential force applied to the petal. Let’s assume that the radius of the flower is R=5 cm, while the tangential force from wind or another source is F=0.1 N. The magnitude of the torque is

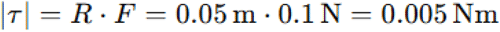

In case of multiple petals (n=5 in *Trachelospermum jasminoides*) experiencing similar forces, the total torque becomes

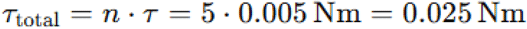

The rotational (*α*) acceleration of the flower petals is related to the torque (Clark and Ryan, 2022) via

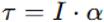

Where I is the moment of inertia of the flower petals about the axis of rotation and α is the angular acceleration. For a flower modeled as a system of point masses at a radius R

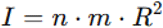

where m is the mass of a single petal. Assuming m=0.002 kg (2 grams per petal):

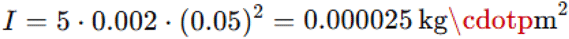

The angular acceleration is:

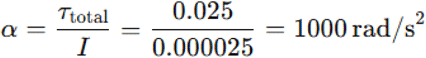

2) The stem experiences forces from bending and torsion, influenced by its helical structure. These forces arise from gravity, wind and the biological tension exerted during growth.

The weight of the stem induces a bending gravitational force. For a stem of length L=20 cm and mass per unit length λ=0.01 kg/m:

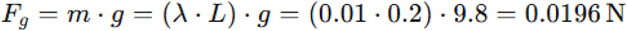

This force acts vertically downward, generating a bending moment at the base of the stem

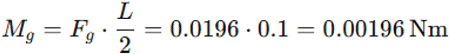

The helical structure of the stem experiences torsional forces due to the winding. The torsional moment (*T*) is given by

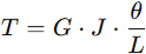

Where G is the shear modulus of the stem material, J is the polar moment of inertia and θ is the angle of twist over the length L.

Assuming G= 10^8^ Pa typical for plant tissue, J = 0.005 m (5 mm radius) and θ = 2π (one full turn over L =0.2 m) (Hoermayer et al., 2024), then

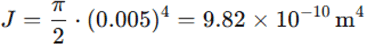

and

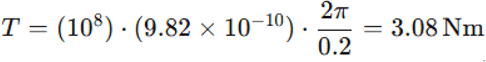

### Internal forces acting within the flower petals and the stem

To calculate the forces within the flower petals and the stem using EST, we need to evaluate the relationship between the macroscopic circulation (observable forces) and the microscopic properties (internal forces or stresses derived from the curl of the force field). The first step is to model the forces using vector fields.

1. Concerning the flower petals, we assume that the external forces (e.g., wind or biological forces) act tangentially to their circular boundary. Further, we assume that the tangential forces induce internal stresses (force per unit area) propagating through the petals. Let the force field acting on the petals be

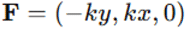

where k is the force constant proportional to the external pressure and x,y represent positions in the z=0 plane.

Next, we compute the curl of the force field. The curl of the force field relates to the internal stresses within the petals. For the flower petals, in the z=0 plane, the curl of **F** = (-*ky, kx*, 0) is

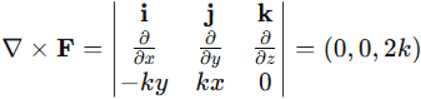

This curl is constant in the z-direction, indicating a uniform internal rotational stress throughout the petals.

2) Concerning the helical stem, it experiences external forces such as gravity (**F**_*g*_) and biological growth forces (**F**_*b*_) that induce internal torsion and bending stresses.

For simplicity, we model the net force field in the stem as: **F** = (−*ky, kx, kz*) where the kz-term accounts for the vertical components of the forces.

Next, we compute the curl of the force field, which provides insight into the internal stresses acting within the stem. This computation reveals the distribution and intensity of these stresses, capturing the complex interplay of forces across the helical structure. For the stem the curl of **F** = (-*ky, kx, kz*) is

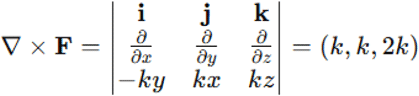

This suggests a complex pattern of internal stress within the stem, with components distributed across all three spatial directions.

#### Visualization and statistics

Diagrams of the flower petals and stem are created to illustrate their geometry, boundary dynamics, and associated vector fields. The Matplotlib library is employed to generate detailed plots, including the circular boundary and tangential forces acting on the flower petals, the curl of the vector field over various surfaces, and the helicoidal path and vector field representation for the stem.

To ensure statistical validation, numerical accuracy is achieved through high-resolution parameterization, with the parameter t sampled at 1,000 points per cycle. The consistency between line integrals and surface integrals is carefully evaluated to confirm the applicability of the extended theorem to the analyzed geometries.

In the sequel, the surface integral of the curl of the vector field will be computed over these surfaces using the extended Stokes’ theorem (EST).

## RESULTS

As stated above, both the flower and the stem experience external and internal mechanical forces that influence their motion and structural behavior:

1. For the flower petals, the torque arising from tangential forces induces a counterclockwise rotation, with the total torque τ_total_ = 0.25Nm and the angular acceleration measured as α=1000 rad/s^2^. The internal stresses in the petals are uniform with a value of 2k and are directly proportional to the external forces acting on them. This proportionality explains the rotational equilibrium observed in the petals.
2. For the stem, the primary forces include a gravitational bending moment *M*_g_ = 0.00196 Nm and a torsional moment T=3.08□ Nm due to a helical twist. The internal stresses in the stem vary in all three dimensions because of its helical geometry. Among these stresses, torsion, proportional to kc, predominates, whereas bending stresses, proportional to kR, have a secondary but still notable influence.

We can now apply EST to relate macroscopic and microscopic circulation. The surface integral of the curl of the vector field is computed over the surfaces, relating the surface integrals to the line integrals along the boundaries. For the flower petals, the counterclockwise macroscopic rotation is calculated by integrating along the circular path in the plane. For the stem, the integral is evaluated over the helical surface.

1. Concerning the flower petals, the boundary of the flower is a circle of radius R. The macroscopic circulation (line integral along the petal boundary) is

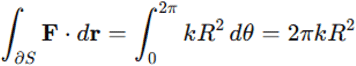

Using the curl, the surface integral is

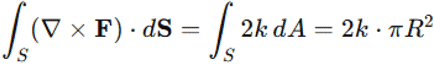

Both results match, confirming that the inner stresses in the petals are proportional to 2k.

In sum, the numerical values for the macroscopic (surface) flows and microscopic (internal) flows in the flower, as governed by EST, are as follows. For the flower, the surface flow (evaluated as a surface integral) is 0.157N\ppm, while the internal flow (evaluated as a line integral) is also 0.157N\ppm. The flower petals exhibit a simple and symmetric geometry, where forces act tangentially along a circular boundary in the z=0 plane. The petals lie on a flat, two-dimensional surface characterized by a constant curl of the force field (∇ ⨯ **F** =(0,0,2*k*)), signifying that the internal forces are uniformly distributed. This uniform distribution creates a direct and proportional relationship between the macroscopic flow (line integral along the circular boundary) and the microscopic flow (surface integral over the disk). The symmetry of the geometry ensures that every contribution to the line integral is exactly matched by the surface integral. Consequently, the uniform geometry and constant curl lead to a perfect agreement between the surface flow and the internal flow, consistent with EST.

2) Concerning the helical stem, the boundary of the stem is parameterized as a helicoidal spiral

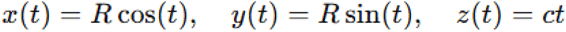

The macroscopic circulation (line integral along the helical path) is

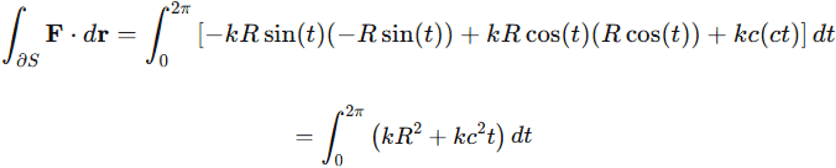

For one turn^(*t* ∈ [0,2π]^

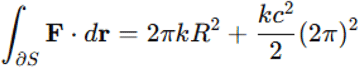

Using the curl, the surface integral is approximated by the ribbon spanned by the helix

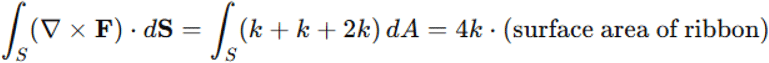

The surface area of the ribbon is

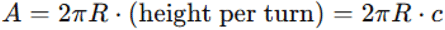

Thus

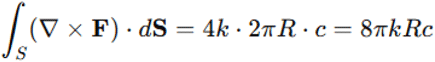

In sum, the numerical values for the macroscopic (surface) flows and microscopic (internal) flows in the stem, as governed by EST, are as follows. For the stem, the surface flow (evaluated as a surface integral) is 2.513N\ppm, while the internal flow (evaluated as a line integral) is 8.053N\ppm. Unlike the flower, the values for surface flow and internal flow differ significantly. This is due to the stem’s more complex geometry, which features a three-dimensional helicoidal structure with a helical boundary and a ribbon-like surface. Unlike the constant curl observed in the flower, ∇ ⨯ **F** = (*k, k*, 2*k*) the curl of the force field in the stem varies in all three dimensions. This non-uniform curl introduces additional contributions to the surface integral that are not directly proportional to the line integral along the helical path. The helicoidal surface spanned by the path is not planar. Its area depends on the radius of the helix and the rise per turn (c), which increases the surface integral significantly compared to the simpler circular geometry of the flower. The line integral along the helical path includes contributions from the vertical rise (z-component), which are absent in the flat geometry of the flower. These vertical components add substantially to the internal flow, making it larger than the surface flow. Forces and circulation in the stem are not confined to a two-dimensional plane, rather display three-dimensional dynamics that capture complex interactions such as bending, twisting and torsional effects, further contributing to the discrepancy between the surface and internal flows. Therefore, the stem’s intricate geometry and three-dimensional dynamics lead to a disparity between surface and internal flows, reflecting the additional factors at play in its structural behavior.

In conclusion,

1. flowers have a circular, symmetric geometry that ensures uniform force distribution and curl. This results in surface and internal flows being equal, as the entire flow field is captured in a flat, two-dimensional setup.
2. In contrast, the stem’s helicoidal geometry introduces non-uniform force distributions and additional components such as vertical contributions and a larger surface area. These factors create a larger internal flow compared to the surface flow, as the line integral accounts for three-dimensional effects that the surface integral does not fully capture.

These differences highlight the impact of geometry and force distribution on the interplay between macroscopic circulation and microscopic forces, showcasing the utility of the extended Stokes’ theorem in analysing forces and circulation in systems exhibiting spiral dynamics.

## CONCLUSION

Classical theorems such as Green’s Theorem (GT) and Stokes’ Theorem (ST) have been pivotal in linking local properties of vector fields to their global behavior. GT applies just to two-dimensional regions and closed curves, while ST extends to three-dimensional spaces requiring closed surfaces or boundaries for its application (Green, 1828; Schey, 1997). These theorems, focused on closed-loop circulations, have proven instrumental in analyzing flows and circulations in systems where boundaries are well-defined, such as steady-state circulations in airflow around wings or electromagnetic field behavior in closed circuits (Arfken, 1985; De Villiers, 2006). However, their utility diminishes when applied to open, three-dimensional trajectories like the helicoidal spirals which are frequently encountered in natural and engineered systems.

We suggest a generalization of ST to establish a mathematical framework connecting the line integral along a helicoidal spiral path to the surface integral of the curl of the vector field over a bounded region. By redefining the boundary concept for helicoidal paths, this framework provides a new tool for analyzing macroscopic and microscopic flow dynamics in complex systems. The EST formulation provides novel insights into the interplay between rotational and translational motions, allowing for a deeper understanding of spiral flows in a variety of physical and biological systems. A key advantage of the extended formulation lies in its ability to model a wide range of systems where spiral or helical dynamics are dominant. For instance, the novel framework enables the analysis of DNA supercoiling, bacterial flagella, biomechanical patterns such as the phyllotaxis of plants (Reinhardt and Gola, 2022; Liu et al., 2024), intracardiac spiral flows observed in cardiac cycles (Mulimani et al., 2022) as well as magnetic vortices in superconductors (Sachkou et al., 2019) and rotational dynamics of spiral galaxies (Blaser et al., 2024), where classical methods fail to capture the intricacies of rotational and translational dynamics.

In this paper, EST is applied to two case studies related with *Trachelospermum jasminoides*, namely the forces acting on flower petals and the helical stress distribution within plant stems.

1. For the flower petals, the circular geometry allows for a straightforward application EST, since the tangential forces acting along the petal boundary produce a uniform curl which is proportional to the rotational stresses. The equivalence between the line integral along the petal boundary and the surface integral of the curl over the enclosed disk validates EST’s effectiveness for two-dimensional spiral systems. The uniform rotational stresses observed in the petals align well with the mathematical predictions of EST. This provides insights into how forces are distributed within the boundary of the flower, potentially aiding in the study of floral mechanics and growth patterns. EST suggests that microscopic forces acting at the level of the petals contribute to the macroscopic rotational motion observed at the flower’s boundary. This could be applied to study the impact of environmental factors like wind on plant structures and to investigate the mechanical interactions between flowers and pollinators during the pollination process. Still, EST effectively simplifies complex calculations by converting a line integral along the flower’s boundary into a surface integral over the petal region. This transformation minimizes computational effort while preserving accuracy.
2. In the case of the stem, although the helical geometry of the stem presents a significant challenge for classical mathematical tools, EST effectively simplifies the intricate interplay of forces involved. The torsional and bending forces are captured through the curl of the vector field, which has components in all three dimensions. The equivalence of the surface integral over the helical ribbon region and the line integral along the helical path demonstrates the robustness of EST in handling three-dimensional geometries with open boundaries. The EST capability to connect macroscopic flow patterns with microscopic circulatory forces may have significant implications for understanding the biomechanics of plant growth and structural stability. This relationship can also provide valuable insights for studies on nutrient and water transport within stems, as these processes often involve spiral dynamics.

Certain assumptions and limitations are inherent in our analysis. EST assumes that the involved vector fields and surfaces are continuously differentiable. In real-world biological systems, irregularities and discontinuities in the geometry or force distribution may reduce the accuracy of the analysis. The flower petals are modeled as a perfect circle and the stem as a regular helix. While this simplifies the mathematical analysis of forces in idealized systems, real-world systems often deviate from these idealized shapes. The analysis of irregular geometries or highly dynamic boundaries may still require significant computational effort, particularly for numerical integration of complex surface and line integrals. The tangential and torsional forces are assumed to be uniform across the boundaries. In reality, biological and environmental forces such as wind, gravity and growth pressures are often spatially and temporally variable. Additionally, secondary effects such as shear forces or anisotropic material properties are not incorporated, which could limit the applicability of the results to certain systems. Future work could extend the framework to handle more irregular and biologically realistic geometries, such as asymmetrical petals or non-uniform stem shapes. The analysis of time-varying forces and boundaries, such as those caused by growth or environmental changes, could provide deeper insights into real dynamics. Integrating the extended theorem with experimental data would help validate the theoretical predictions and refine the mathematical models.

In conclusion, the proposed extension to Stokes’ Theorem integrates helicoidal paths into circulation analysis, bridging a critical gap and expanding its applicability to open, non-planar trajectories. By redefining boundaries, it simplifies the study of rotational and translational flows, offering a versatile tool for analyzing complex dynamics such as those observed in the flowers and stem of *Trachelospermum jasminoides*.

## DECLARATIONS

## Ethics approval and consent to participate

This research does not contain any studies with human participants or animals performed by the Author.

## Consent for publication

The Author transfers all copyright ownership, in the event the work is published. The undersigned author warrants that the article is original, does not infringe on any copyright or other proprietary right of any third part, is not under consideration by another journal, and has not been previously published.

## Availability of data and materials

all data and materials generated or analyzed during this study are included in the manuscript. The Author had full access to all the data in the study and take responsibility for the integrity of the data and the accuracy of the data analysis.

## Authors’ contributions

The Author performed: study concept and design, acquisition of data, analysis and interpretation of data, drafting of the manuscript, critical revision of the manuscript for important intellectual content, statistical analysis, obtained funding, administrative, technical, and material support, study supervision.

## Declaration of generative AI and AI-assisted technologies in the writing process

During the preparation of this work, the author used ChatGPT to assist with data analysis and manuscript drafting. After using this tool, the author reviewed and edited the content as needed and takes full responsibility for the content of the publication.

## Acknowledgements

none.

